# Identifying novel chemical matter against the Chikungunya virus nsP3 macrodomain through crystallographic fragment screening

**DOI:** 10.1101/2024.08.23.609196

**Authors:** Jasmin C. Aschenbrenner, Andre Schutzer de Godoy, Michael Fairhead, Charles W.E. Tomlinson, Max Winokan, Blake H. Balcomb, Eda Capkin, Anu V. Chandran, Mathew Golding, Lizbe Koekemoer, Ryan M. Lithgo, Peter G. Marples, Xiaomin Ni, Warren Thompson, Conor Wild, Mary-Ann E. Xavier, Daren Fearon, Frank von Delft

## Abstract

Chikungunya virus (CHIKV) causes severe fever, rash and debilitating joint pain that can last for months ^1,2^or even years. Millions of people have been infected with CHIKV, mostly in low and middle-income countries, and the virus continues to spread into new areas due to the geographical expansion of its mosquito hosts. Its genome encodes a macrodomain, which functions as an ADP-ribosyl hydrolase, removing ADPr from viral and host-cell proteins interfering with the innate immune response. Mutational studies have shown that the CHIKV nsP3 macrodomain is necessary for viral replication, making it a potential target for the development of antiviral therapeutics. We, therefore, performed a high-throughput crystallographic fragment screen against the CHIKV nsP3 macrodomain, yielding 109 fragment hits covering the ADPr-binding site and two adjacent subsites that are absent in the homologous macrodomain of SARS-CoV-2 but may be present in other alphaviruses, such as Venezuelan equine encephalitis virus (VEEV) and eastern equine encephalitis virus (EEEV). Finally, a subset of overlapping fragments was used to manually design three fragment merges covering the adenine and oxyanion subsites. The rich dataset of chemical matter and structural information discovered from this fragment screen is publicly available and can be used as a starting point for developing a CHIKV nsP3 macrodomain inhibitor.

## Introduction

Chikungunya virus (CHIKV) is a mosquito-borne alphavirus.^1^ Transmitted in urban areas by *Aedes aegypti* and *Aedes albopictus* mosquitos, it poses a significant health threat as seen by recent outbreaks driven by the spread of the mosquito vectors to additional geographical areas.^2^ First discovered in Tanzania in the 1950s^3,4^, it has caused numerous outbreaks since 2004 resulting in millions of infections as it has spread from tropical into subtropical areas and the Western hemisphere.^5^ Patients infected with CHIKV present with severe fever, rash and debilitating joint pain.^6^ In up to 60 % of patients, the joint pain will persist for months or even years after the initial infection, severely impacting the patient’s quality of life.^7^ Currently, there is only one FDA-approved vaccine (IXCHIQ) for protecting adults at risk of exposure to CHIKV available.^8^ Valneva, the company producing IXCHIQ, will first focus on selling the vaccine to travellers who plan to visit countries where the disease is endemic.^9^ To make the vaccine accessible to low and middle-income countries (LMIC) that are hardest hit by CHIKV, Valneva has signed an agreement with Instituto Butantan in Brazil which will develop, manufacture, and market the vaccine in LMIC which is supported by funding from the European Union’s Horizon 2020 programme.^10^ No antivirals for the prevention or treatment of CHIKV are available at this point.^11^

CHIKV is an enveloped, single-stranded positive genomic RNA virus.^1^ Its genome encodes two open reading frames (ORFs) for four non-structural (nsP1, nsP2, nsP3, nsP4) and three structural proteins (capsid and two envelope proteins with two cleavage products) as polyproteins, respectively.^12-14^ The non-structural proteins are involved in transcription and necessary for CHIKV replication.^15^ The nsP1 protein containing methyltransferase and guanylyltransferase domains is involved in viral RNA capping and mediates the assembly of the replication complex.^16,17^ The nsP2 of CHIKV has RNA-modulating activity via its helicase, nucleoside triphosphatase, RNA-dependent 5’-triphosphatase, and protease.^18-20^ The C-terminal cysteine protease of nsP2 processes the non-structural polyprotein (P1234) into the individually active nsPs, and has been the focus of drug discovery efforts due to its importance in the viral replication cycle.^21-23^ nsp4 contains the RNA-dependent RNA polymerase of CHIKV which produces negative-sense, genomic and sub-genomic viral RNAs performing both primed extension and terminal adenylyltransferase activities.^24^ Both proteases and polymerases are well-validated antiviral targets, as demonstrated by the development of successful therapeutics for HIV, hepatitis, and SARS-CoV-2.^25-29^ Therefore, these could be considered lower-risk projects compared to targeting macrodomains.

nsP3 forms part of the replicase unit and functions as a stand-alone protein.^30^ It can be found both in membrane-bound replication complexes and large cytoplasmic non-membranous granules.^31^ nsP3 consists of three distinct domains: an N-terminal macrodomain, a central zinc-binding domain called the alphavirus unique domain (AUD) and the C-terminal poorly conserved and highly phosphorylated hypervariable domain (HVD).^30^ The nsP3 macrodomain is highly conserved between alphaviruses and homologs can be found in other positive-strand RNA viruses, such as rubella virus, some coronaviruses and hepatitis E virus.^32-35^ It functions as an ADP-ribosyl (ADPr) hydrolase, removing mono-ADPr from post-translationally modified aspartate and glutamate residues of viral and host-cell proteins, counteracting the innate immune response.^31,32,36-38^ Deficiency in the ADPr-binding and hydrolase function of the nsP3 macrodomain was shown to attenuate CHIKV virulence in mice and to lead to reduced viral replication in cell culture, making it a potentially attractive target for antiviral research.^31^

Crystallographic fragment screening is a well-validated method for the identification of novel chemical matter for the development of potent lead molecules.^39,40^ Hit identification using X-ray crystallography has the advantage of providing detailed information about the specific protein-ligand interactions and protein dynamics. Furthermore, the obtained structural information can accelerate subsequent optimisation efforts.^41^ In this study, we performed a high-throughput, crystallographic fragment screen on the nsP3 macrodomain of CHIKV. We screened a total of 1385 fragments, yielding 109 fragment hits covering the ADPr-binding site of CHIKV nsP3 macrodomain and adjacent subsites. Our openly available structural data aims to provide valuable information for the design of follow-up compounds with higher affinity for the CHIKV nsP3 macrodomain.

## Materials and methods

### Cloning of Chikungunya macrodomain of nsP3

The coding region of the CHIKV macrodomain (Genbank ID ACD93569.1) from the Chikungunya virus strain S27-African prototype (Taxonomy ID: 371094), corresponding to residues 1334 to 1493 of the full-length polyprotein, was codon optimized for *E. coli* expression and synthesized by Genscript. The encoding region was PCR-amplified and inserted into pETM11/LIC by ligation-independent cloning (LIC)^42^. Finally, the region encoding the CHIKV macrodomain was cloned into the pET28 vector with a TEV cleavable hexahistidine tag using Gibson assembly.

### Expression and purification

Expression and purification were performed using the PREPX workflow^43^, described below. Typically, 100 ng of plasmid was transformed into the *E. coli* protein expression strain BL21(DE3), and a single colony was used to inoculate 10 mL of SOC containing 50 µg/mL kanamycin. The starter culture was grown overnight at 37°C 250 rpm shaking and was used the following day to inoculate 1 L of TB (ForMedium) in a 2.5 L baffled shake flask supplemented with 50 µg/mL kanamycin and 0.01% Antifoam-204 (Merck). Cells were grown first for 3 h at 37°C and then for 2 h at 18°C, 250 rpm shaking. Protein expression was then induced with 0.5 mM Isopropyl-β-D-thiogalactopyranosid (IPTG), and growth continued overnight at 18°C, 250 rpm shaking. Cells were harvested by centrifugation at 4,000x*g* and the pellets were re-suspended using 3 mL of Base Buffer (10 mM HEPES, 500 mM NaCl, 5 % glycerol, 30 mM imidazole, 0.5 mM TCEP, pH 7.5) per gram wet weight of cells. Triton X-100, lysozyme and benzonase to a final concentration of 1 %, 0.5 mg/mL and 1 µg/mL, respectively, were then added before freezing at -80°C. The cell lysate mixture was first thawed in a room-temperature water bath and then centrifuged at 4800x*g* for 1 h at 4°C. The supernatant (soluble fraction) was then loaded onto a 1 mL His GraviTrap column (Cytiva) and washed twice with 20 mL of Base Buffer. Four His GraviTrap columns were used for every 1 L of culture originally grown. The tagged-target protein was then eluted using 2.5 mL of Base Buffer containing 500 mM imidazole and immediately desalted using a PD-10 column (Cytiva) with Base Buffer as the mobile phase. The His tag was then removed by protease digest overnight at 4°C using 1 mg of TEV per 10 mg of target protein. The tag, un-cleaved protein, and TEV (bearing a non-cleavable his-tag) were then removed by passing through a 1 mL His GraviTrap column. The cleaved protein sample was then injected onto a superose 12 pg SEC column (Cytiva) using 25 mM Tris, 100 mM NaCl, 5 % glycerol, pH 7.5, as the mobile phase. Peak nsP3 macrodomain fractions were concentrated to 11 mg/mL before flash freezing in liquid nitrogen and storage at -80°C. Efficacy of the process and final protein purity were assessed by SDS-PAGE using NuPAGE 4-12 % Bis-Tris Midi Protein Gels (ThermoFisher Scientific).

### Crystallisation and fragment screen

Initial conditions for crystallisation of CHIKV Mac were found in the BCS Screen^44^ (Molecular Dimensions). After optimisation, crystals grew reproducibly using sitting-drop vapour diffusion in MRC 3 Lens Crystallisation plates (SWISSCI) with a reservoir solution containing 0.1 M Tris, pH 7.8, 0.1 M Potassium thiocyanate, 0.1 M Sodium bromide, and 25 % (v/v) PEG Smear Broad with drops of 75 nL CHIKV Mac at 11 mg/mL, 40 nL of reservoir solution and 35 nL of CHIKV Mac crystal seed solution.^45^

We screened 1385 fragments from the DSi-Poised Library^46^, Minifrags^47^, Fraglites^48^, Peplites^49^, York 3D^50^, SpotXplorer^51^, and Essential Fragment Library (Enamine). Fragments were soaked into drops containing CHIKV Mac crystals at 25 % (v/v) DMSO using acoustic dispensing^52^ with an Echo 650 liquid handler (Labcyte). After incubation for 1-3 h at 20°C, crystals were harvested and cryo-cooled in liquid nitrogen without additional cryoprotectant.

Data was collected at the I04-1 beamline (Diamond Light Source, UK) at 100 K and automatically processed with Diamond Light Source’s autoprocessing pipelines using XDS^53^ and either xia2^54^, autoPROC^55^ or DIALS^56^ with the default settings.

Analysis of fragment binding was done using the XChemExplorer^57^. Electron density maps were generated with DIMPLE^58^, ligand restraints were calculated with GRADE^59^ and ligand-binding events were identified using PanDDA2^60^. Ligands were modelled into PanDDA2-calculated event maps using its autobuild function^60^ or manually using Coot^61^, and structures were refined by ensemble-refinement of ligand-free and ligand-bound states^62^ using Refmac^63^.

Coordinates, structure factors and PanDDA2 event maps for the ground-state and ligand-bound CHIKV Mac structures discussed here are deposited in the Protein Data Bank (group deposition G_1002294). Data collection and refinement statistics are summarised in Supplementary Data File 1.

### Fragment merges

Datasets were aligned by XChemAlign^64^ before importing into SeeSAR 13.0.1, which was used to design fragment merges manually.

## Results

### Fragment-binding to the nsP3 macrodomain of CHIKV

The CHIKV nsP3 macrodomain is a mono-ADP-ribosyl hydrolase which efficiently binds mono-ADPr and cleaves it from post-translationally modified proteins. The globular protein consists of six β-sheets and five α-helices with an extensive binding pocket to accommodate the substrate. The binding cavity can be divided into smaller sites here named the adenine site, phosphate site, distal ribose site and oxyanion hole, according to the ADPr feature bound in the site (Figure 1).

**Figure 1.**
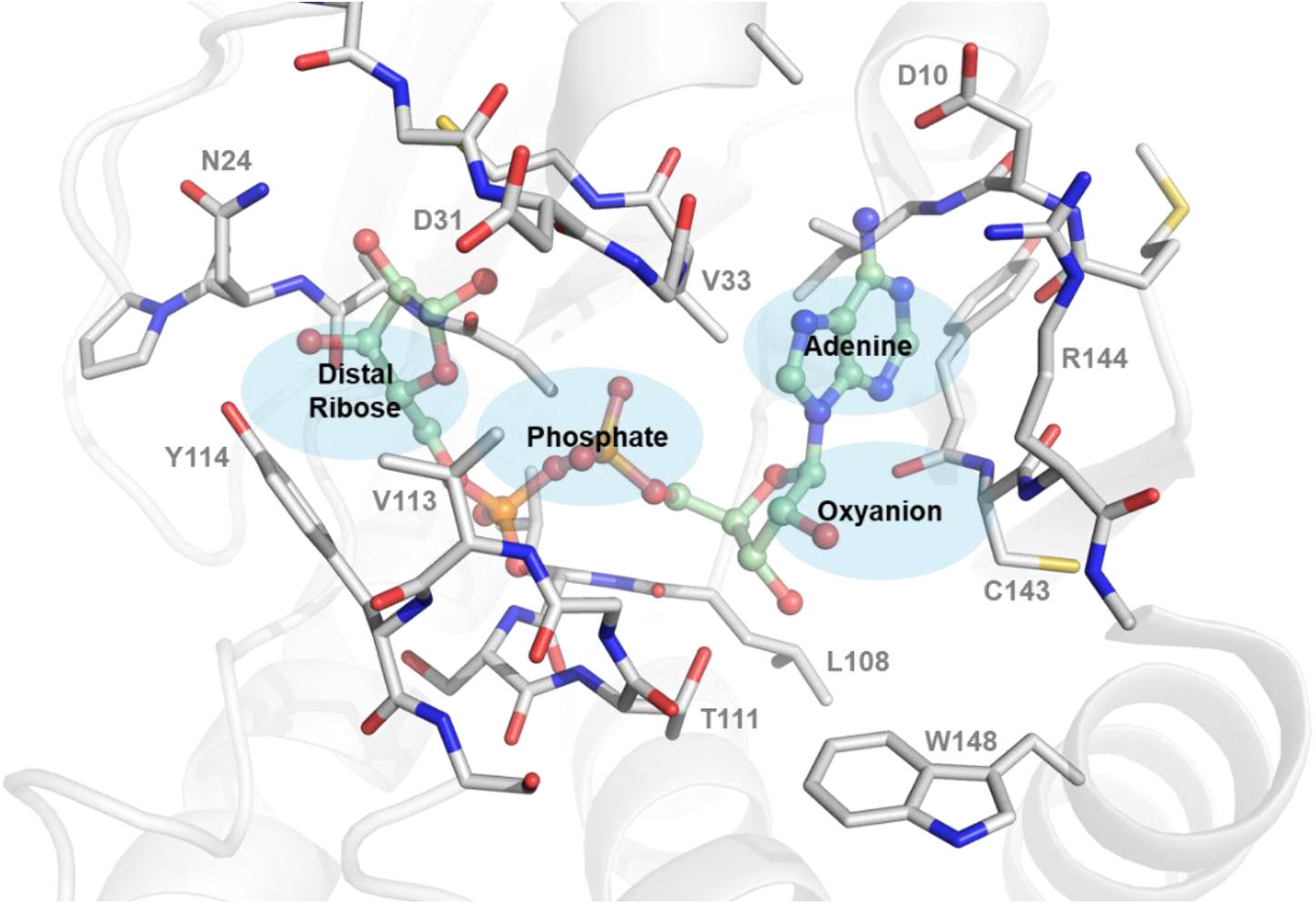
Structure of the ADP-ribose bound nsP3 macrodomain of CHIKV. The ADPr binding site of CHIKV nsP3 macrodomain can be divided into smaller pockets according to the chemical groups and interactions found between ADPr and macrodomain into adenine site, oxyanion site, phosphate site and the (distal) ribose site, which is the catalytic site. The CHIKV nsP3 macrodomain structure (6VUQ) with bound ADPr (pale green) is shown in grey. Residues within 4 Å to the bound substrate are shown as sticks.

Several residues of the CHIKV nsP3 macrodomain are involved in the binding of ADPr: Binding of the adenine is stabilised by hydrogen bonds formed with D10 and R144. Alternatively, cation-π stacking can sometimes be observed between R144 and the pyrimidine of the adenine. In other macrodomains, such as SARS-CoV-2 nsp3 macrodomain and human MacroD2, the adenine moiety undergoes π-π stacking with phenylalanine which is replaced by R144 in CHIKV.^34^ The proximal ribose in the oxyanion hole forms multiple water bridges with the amides of the protein backbone of R144 and L108. Additionally, the hydroxyl group of the ribose can form hydrogen bonds with T111 and the carbonyl of the backbone. The phosphate site is enclosed by two loops that interact via multiple amide-mediated hydrogen bonds with the pyrophosphate of the ADPr. The distal ribose is coordinated via hydrogen bonds to N24 and can form additional water bridges to Y114, its pyrophosphate moiety and the surrounding protein backbones.

Mutational studies have previously attempted to determine key residues involved in substrate binding and catalysis of the CHIKV nsP3 macrodomain. Using virtual alanine scanning and site-directed mutagenesis, it was shown that mutation of N24 or Y114, both close to the distal ribose, or V33, close to the adenosine moiety, to an alanine residue, CHIKV nsP3 macrodomain exhibited lower hydrolase activity, possibly due to affected substrate recognition and binding.^65^ Another study investigated residues D10, G32, T111, G112, and R144. Interestingly, the R144A mutant exhibited similar activity to the wildtype macrodomain and when transfected into mammalian cells as RNA for virus production, could not be rescued as it had reverted to the wildtype residue.^31^ Mutants D10A, G32E and G112E did not exhibit detectable ADPr binding, and their catalytic activity was reduced to negligible levels. Similarly, for R144A, these mutants could not be rescued as the viruses produced from transfection did not carry the mutation.^31^ Mutational studies provide therefore excellent information on key residues and mutational variability that should be used to guide medicinal chemistry efforts during drug discovery.

For our high-throughput, crystallographic fragment screen campaign, we soaked 1385 fragments into the P31 crystal form of CHIKV nsP3 macrodomain with 4 copies in the ASU. The average resolution of the 1082 successfully collected and processed datasets was 1.5 Å. We found a total of 109 fragments binding to the ADPr-binding site or in closely located subsites (Figure 2A). Some fragments, particularly from the Minifrags library^47^, bound multiple times to the large cavity of the ADPr-binding site. Similar to the fragment screens of the nsp3 macrodomain of SARS-CoV-2^66^, the majority of fragment hits were identified in the adenine site (62 hits). Only a few fragments (9) were found binding to the phosphate site, but this was more chemical matter than was found against SARS-CoV-2 Mac1 with only a single fragment hit in this site^66^. For the sites occupied by the terminal ribose and the oxyanion subsite, 15 and 33 fragments were observed, respectively.

**Figure 2.**
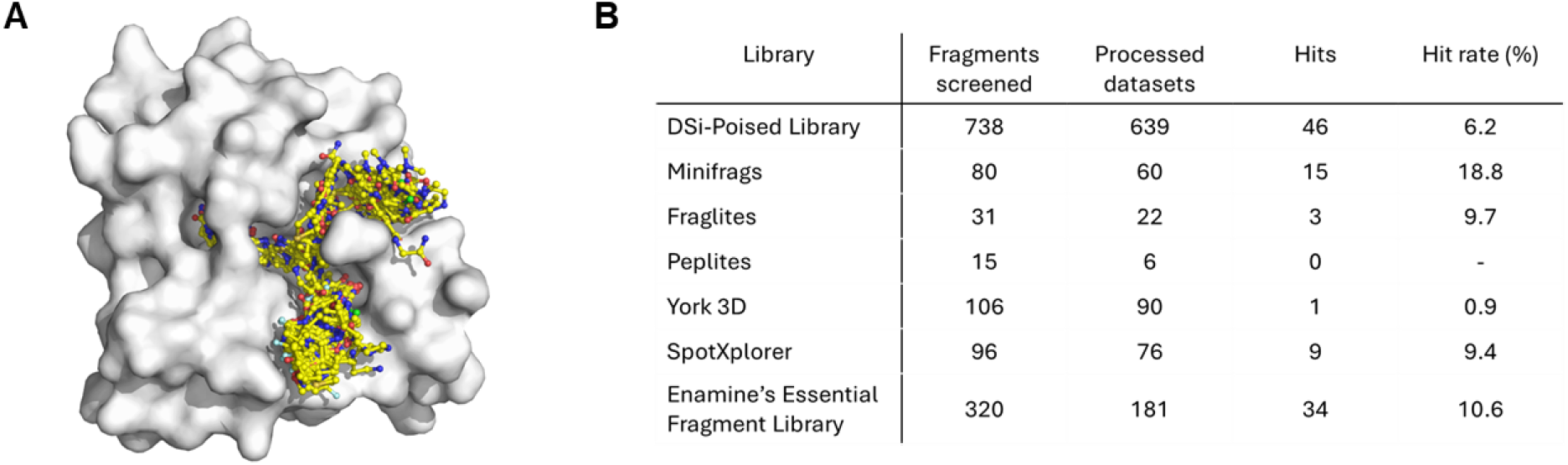
Crystallographic fragment screening of the CHIKV nsP3 macrodomain identified 109 fragments bound to the ADPr binding site and nearby subsites. (**A**) Overview of fragments (yellow) bound to ADPr binding site and nearby subsites on the CHIKV nsP3 macrodomain (shown as surface representation). (**B**) Overview of the fragments screened, the number of processed datasets, the hits found from each library and the calculated hit rate (%). ‘Processed datasets’ refers to the datasets that could successfully be solved by Dimple^58^ and were subsequently analysed by PanDDA2^60^. From the 1385 fragments screened, we obtained 1074 processable datasets. 311 datasets (22 %) were not obtained or analysed due to crystal pathologies.

### Analysis of key interactions of fragments binding to CHIKV nsP3 macrodomain

#### Adenine site

A total of 62 fragments were found to bind in the adenine site of CHIKV nsP3 macrodomain. Most could be classified into one of six commonly found scaffolds: amino benzimidazoles/benzothiazoles, aminoquinolines/benzoxazoles, pyrrolopyridines, aminopyridine and aminopyridine-like heterocycles, amides or benzodioxoles/-dioxanes/-furans (Figure 3).

**Figure 3.**
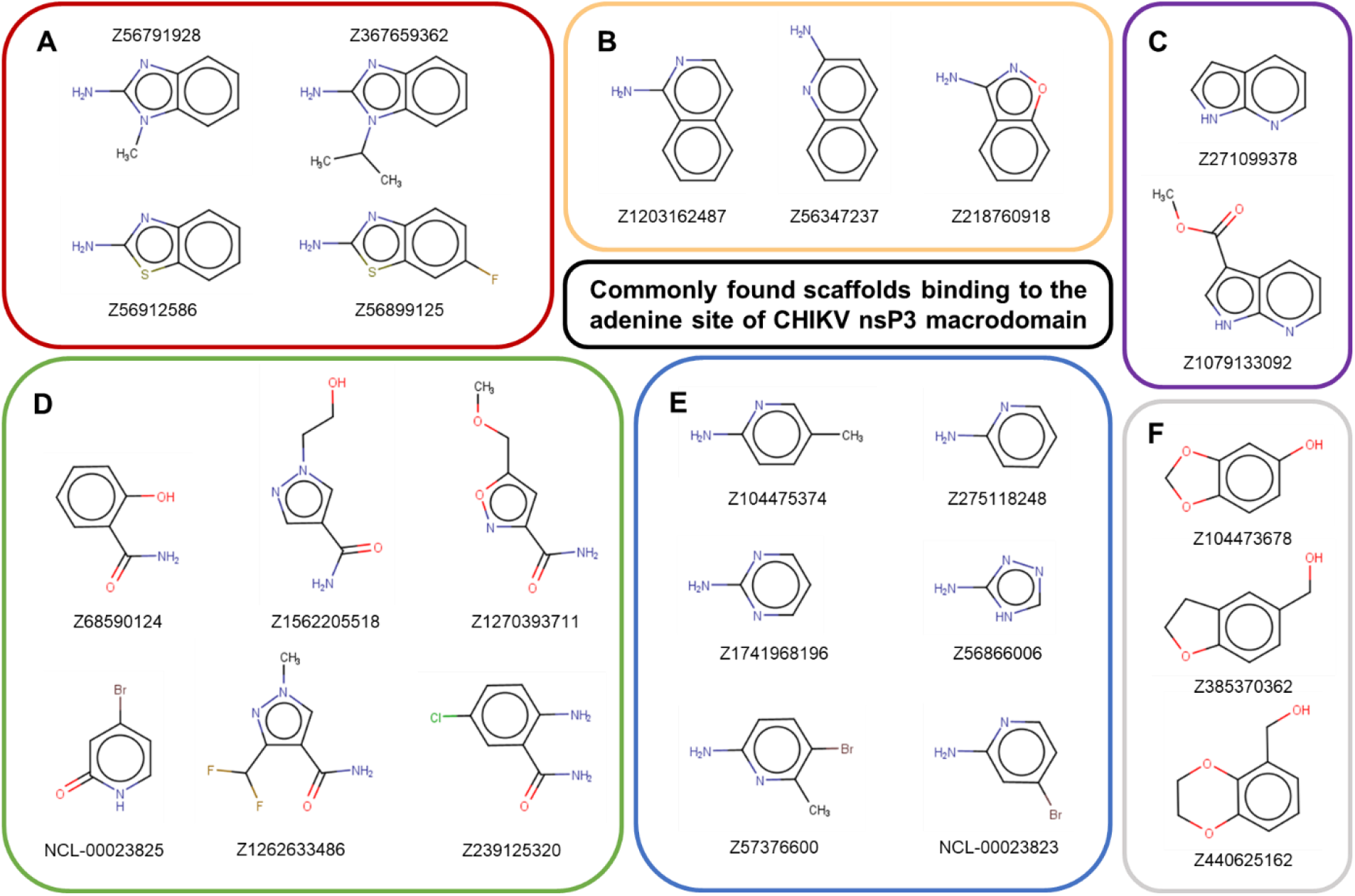
Common scaffolds found in fragments binding to the adenine site of CHIKV nsP3 macrodomain. The majority of the fragments found binding to the adenine site could be grouped into (A) amino benzimidazoles/benzothiazoles, (B) aminoquinolines/benzoxazoles, (C) pyrrolopyridines, (D) amides, (E) aminopyridine-like, (F) benzodioxoles/-dioxanes/-furans.

A common motif in the observed fragments from the amino benzimidazoles/benzothiazoles, aminoquinolines/benzoxazoles, pyrrolopyridines, and aminopyridine-likes is the 1,3-hydrogen bond donor/acceptor motif forming interactions with the side-chain oxygen of Asp10 and amide of Ile11, similar to the adenosine moiety of ADPr (Figures 4A, B_iv-x_). This interaction resembles the hinge-binding motif commonly found in protein kinase inhibitors.^67^ The same interactions were found with fragment scaffolds containing an amide group, forming hydrogen bonds to Asp10 and Ile11 (Figure 4B_i-iii_) with some showing additional interactions, such as water bridges to the carbonyl group of Met9 (Z1562205518, Figure 4B_i_) or a hydrogen bond to the carbonyl of Tyr142 via an amine group (Z239125320, Figure 4B_ii_). More distinct to the nitrogen-containing scaffolds was the group of benzodioxoles/-dioxanes/-furans where the interaction was dominated by a single hydrogen bond from the oxygen of the fragment to the amide bond of Ile11 as seen with Z104473678 (Figure 4B_xi_).

**Figure 4.**
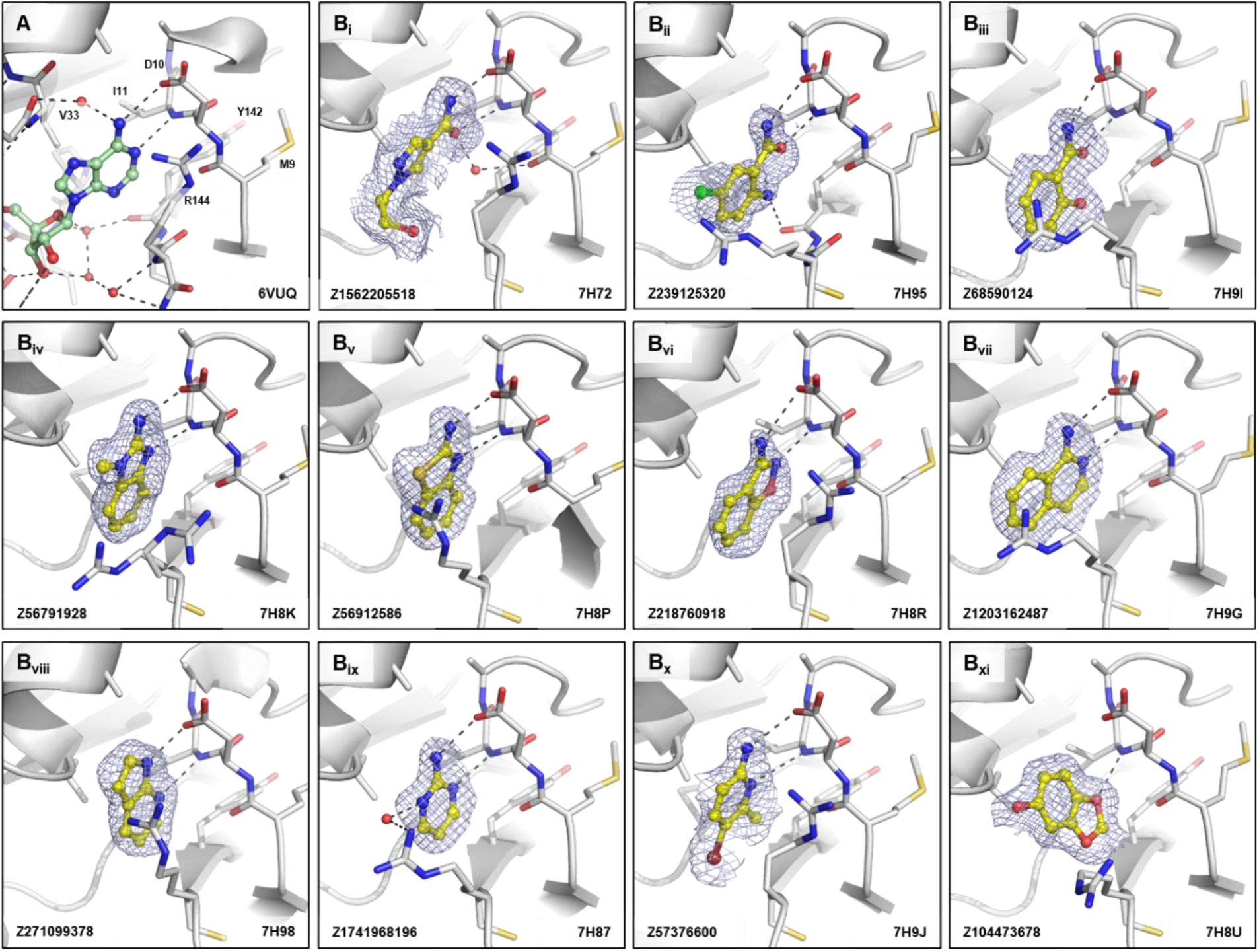
Fragments binding to the adenine site of CHIKV nsP3 macrodomain. (**A**) Stick representation of ADPr (pale green) binding to the adenine site. (**B**_**i-xi**_) Examples of representative fragments bound to the adenine site in CHIKV nsP3 macrodomain. The CHIKV nsP3 macrodomain structure is shown in grey. The bound fragment is shown with yellow sticks surrounded by a blue mesh generated from the PanDDA2 event map. Polar interactions are highlighted by black dashes.

#### Oxyanion site

Lying below the adenine subsite is the oxyanion hole which is characterised by interactions with the backbone nitrogens of Arg144 and Asp145. If ADPr is bound to the CHIKV nsP3 macrodomain, a distinct water network is visible connecting the proximal ribose of ADPr with the amide bonds of Leu108, Arg144 and Asp145, as well as the carbonyl of Tyr142 by water bridges (Figure 5A). Fragments found to bind in the oxyanion hole generally feature a carbonyl pointing towards the same backbone nitrogens, forming hydrogen bonds with one or both nitrogens (Figure 5B). While the crystallographic fragment screen of the SARS-CoV-2 nsp3 macrodomain revealed chemically and geometrically diverse hits in the oxyanion site with interactions formed by carboxylates, sulfones, isoxazole, α-keto acid, and a succinimide^66^, the fragments binding at the oxyanion subsite in CHIKV nsP3 macrodomain feature mainly heterocycles such as pyridones, piperidones, pyridazinones, and piperazinones, likely influenced by Trp148 which offers π-stacking interactions such as seen with Z3219959731 (7H89), Z1162778919 (7H6S), and Z1154747269 (7H6R) (Figure 5B_i,iv,v_).

**Figure 5.**
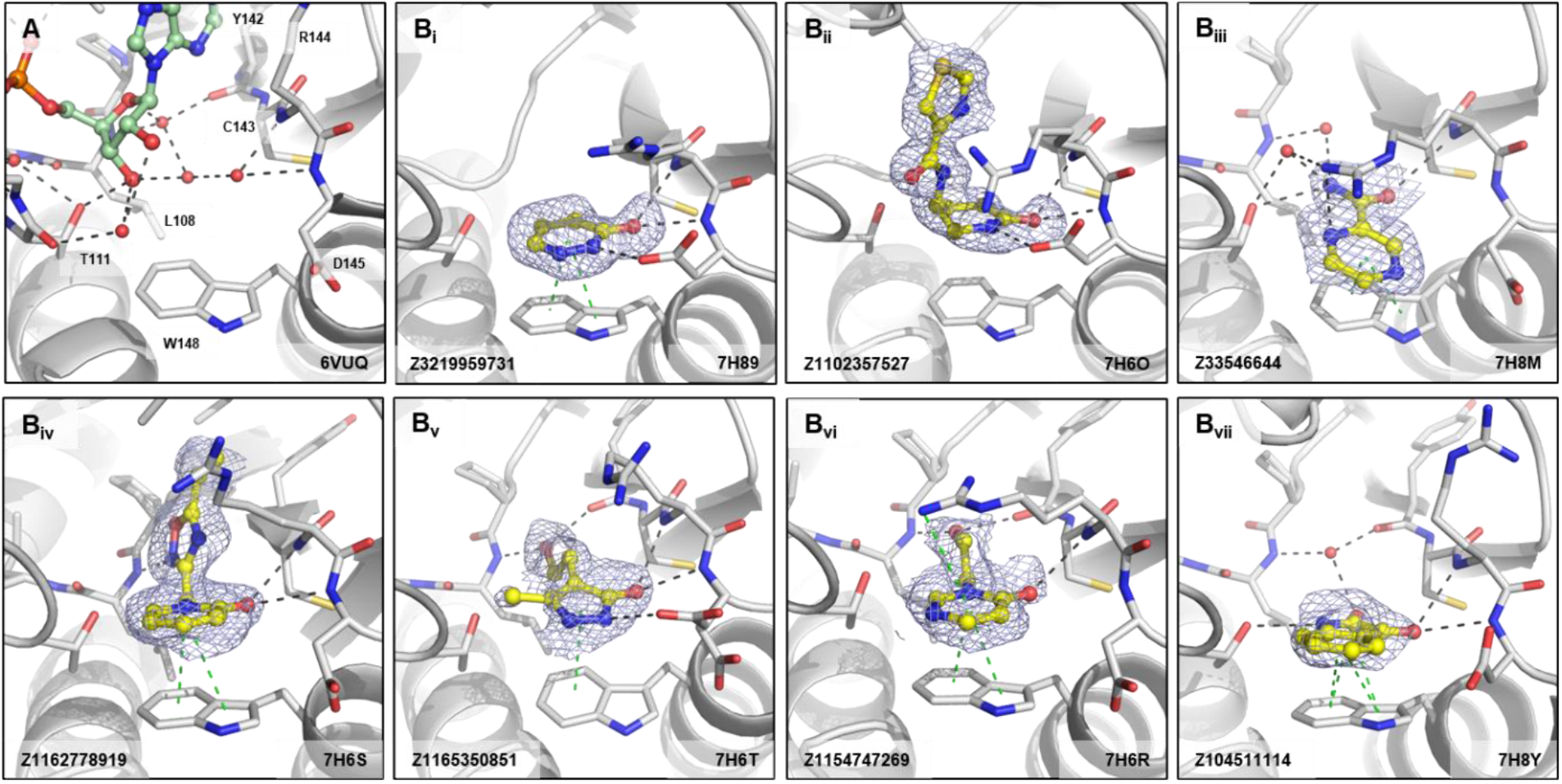
Fragments binding to oxyanion site of CHIKV nsP3 macrodomain. (**A**) Stick representation of ADPr (pale green) binding to the CHIKV nsP3 macrodomain with view of the water network mediating polar interactions to the proximal ribose of ADPr. (**B**_**i-vi**_) Examples of representative fragments bound to the oxyanion site in CHIKV nsP3 macrodomain. The CHIKV nsP3 macrodomain structure is shown in grey. The bound fragment is shown with yellow sticks surrounded by a blue mesh generated from the PanDDA2 event map. Polar interactions and π-stacking interactions are highlighted by black and green dashes, respectively.

#### Ribose-binding and Phosphate-binding site

Only a small percentage (13 %) of the bound fragments were found at the ribose-or phosphate-binding subsites with most of them being extremely low molecular weight fragments, often a single heterocycle (9 out of 14 fragments). No fragment hits were observed on the ‘outer’ side of Tyr114 close to Asn24 and Pro25 in the CHIKV nsP3 macrodomain in contrast to the crystallographic fragment screen on the SARS-CoV-2 nsp3 macrodomain^66^, suggesting that those fragments may be an artefact of crystal packing.

In the phosphate binding site, the diphosphate group of ADPr forms hydrogen bond interactions with protein backbone amides or carbonyls of residues Ala22, Val33, Cys34, Leu108, and Ser110-Tyr114 either directly or mediated via two highly coordinated water molecules (Figure 6A). Similar interactions were exploited by the fragments, with interactions in the phosphate tunnel mostly based on hydrogen bonds between the fragments and the protein backbone, sometimes mediated by water molecules (Figure 6B_i-v_). A common feature found was a water bridge from an aromatic nitrogen atom of the fragments to a water connecting to the amide of Val113, such as in Z1741970824 (Figure 6B_v_). An interesting observation was made with Z1041785508. In addition to forming hydrogen bonds, the triazolyl-thiazole scaffold interacts with Tyr114 via π-stacking and induces a peptide flip of Thr111, potentially due to π-stacking and hydrophobic interactions (Figure 6B_i_).

**Figure 6.**
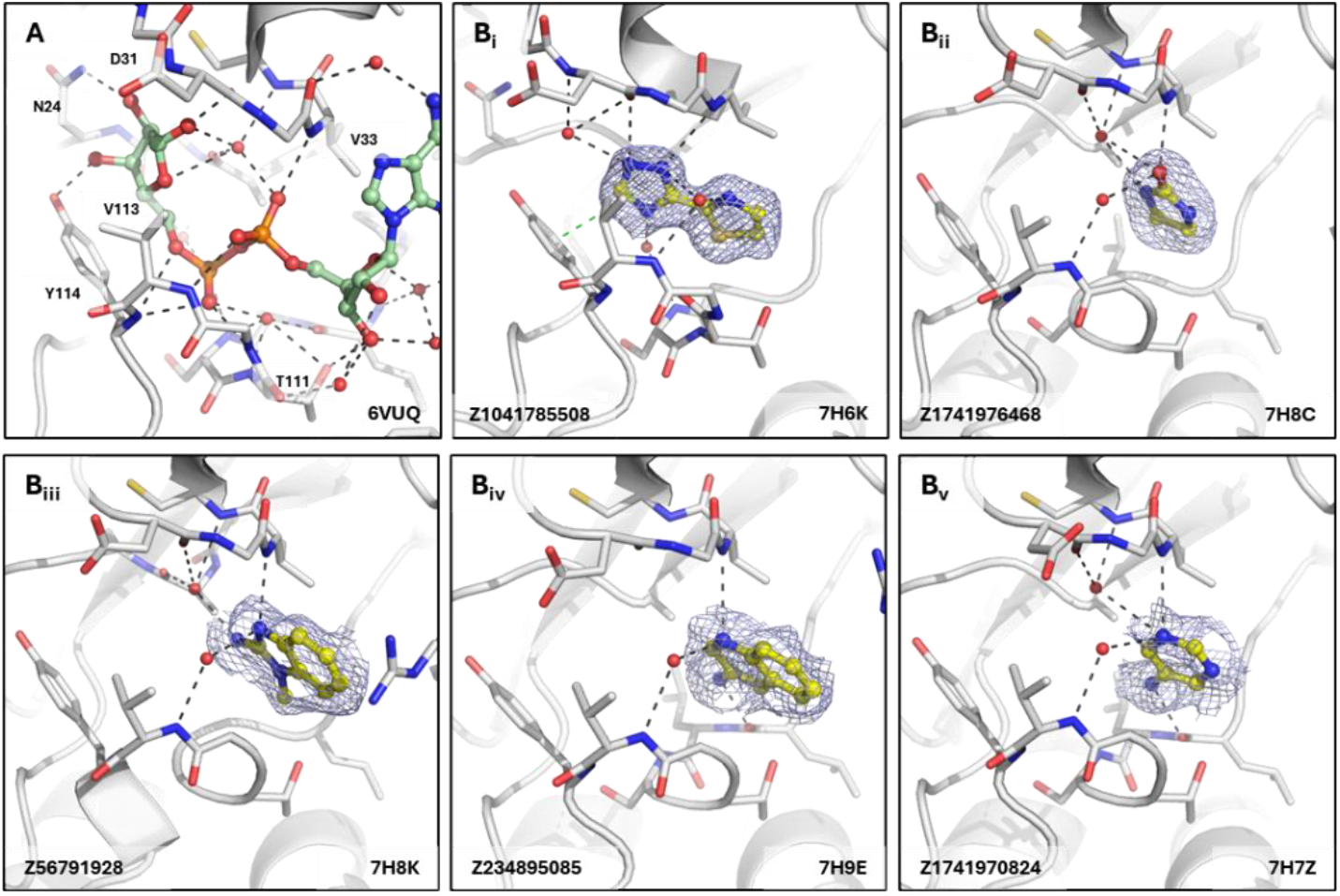
Fragments binding to phosphate site of CHIKV nsP3 macrodomain. (**A**) Stick representation of ADPr (pale green) binding to the CHIKV nsP3 macrodomain with view of the water network mediating polar interactions to the diphosphate and the ribose of the adenosine. (**B**_**i-vi**_) Examples of representative fragments bound to the phosphate site in CHIKV nsP3 macrodomain. The CHIKV nsP3 macrodomain structure is shown in grey. The bound fragment is shown with yellow sticks surrounded by a blue mesh generated from the PanDDA2 event map. Polar interactions and π-stacking interactions are highlighted by black and green dashes, respectively.

The distal ribose-binding site, which is the catalytic site, coordinates the distal ribose of the bound ADPr via hydrogen bond interactions of the hydroxyl groups to Asn24 and Tyr114. The oxygen in the ribose of ADPr forms a water bridge to a highly coordinated water interacting with the carbonyl of Ala22 and the amide of Cys35 (Figure 7A). Fragments, that were found to bind in the ribose site, contained either a carbonyl (as in the pyrrolid(in)one scaffolds Z57127349 and Z940713508 and the pyridazinone scaffold Z3219959731), a sulfone as in Z303754556, or a nitrogen group (Z1154747269), interacting with the Asn24 sidechain and the backbone amide of Asp31 via by hydrogen bonds (Figure 7B_i-vii_). The second carbonyl of the pyrrolidinone scaffold Z57127349 forms water-mediated polar interactions with the amide of Val33 and, interestingly, the backbone carbonyl of Thr111 which exhibits a peptide flip similarly to when the triazolyl-thiazole fragment (Z1041785508) binds to the phosphate site (Figure 7B_i_ and Figure 6B_i_). In addition to polar interactions, some of the mini-frags display π-stacking interactions to Tyr114, such as seen with Z1741976468 (Figure B_iv_) and Z1741970824 (Figure B_v_).

**Figure 7.**
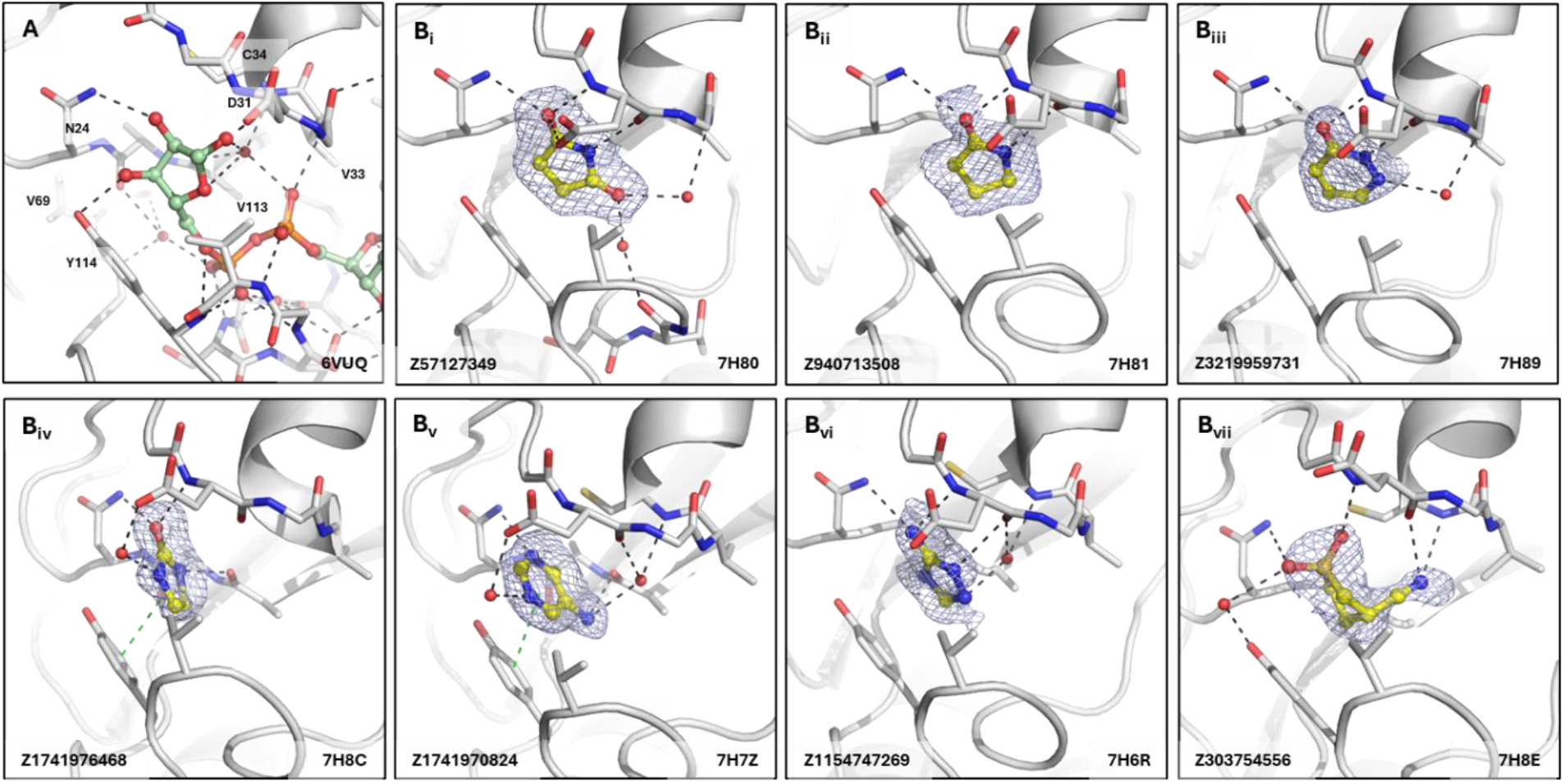
Fragments binding to ribose site of CHIKV nsP3 macrodomain. (**A**) Stick representation of ADPr (pale green) binding to the CHIKV nsP3 macrodomain with view of the water network mediating polar interactions to the distal ribose of the ADPr. (**B**_**i-vi**_) Examples of representative fragments bound to the ribose site in CHIKV nsP3 macrodomain. The CHIKV nsP3 macrodomain structure is shown in grey. The bound fragment is shown with yellow sticks surrounded by a blue mesh generated from the PanDDA2 event map. Polar interactions and π-stacking interactions are highlighted by black and green dashes, respectively.

A virtual screening effort against the CHIKV nsP3 macrodomain by Zhang et al. previously yielded co-crystal structures with 8 different pyrimidone scaffolds, all binding to the ribose site.^68^ Interestingly, the pyrimidone fragments displayed a slightly different binding mode in comparison to what we observed in our study. The pyrimidone-carboxylic acid fragments, e.g. in 6W8Y, bound parallel to Tyr114, forming π-stacking interactions, but with the carbonyl pointing towards Gly70, which is part of the loop making up the ‘backside’ of the pocket, and the carboxy group forming hydrogen bonds to the backbone nitrogen of Tyr114 and Gly112. This was not observed in our fragment hits, which bound further up and closer to Asn24 and Asp31 in the ribose site.

#### Fragments identified in other subsites

Our crystallographic fragment screen revealed fragment binding in two sub-sites distinct to those directly involved in substrate recognition. 25 fragments revealed a cryptic pocket between Arg144 and Met9/Asp10. This pocket, designated by us as the ‘Arg144 subsite’, was not present in the ADPr-bound CHIKV nsP3 macrodomain structure (Figure 8A) and is only accessible by certain fragments due to the flexibility of the Arg144 sidechain. The equivalent residue in the SARS-CoV-2 macrodomain, Phe156, has extremely limited conformational flexibility, making this subsite unique to the CHIKV nsP3 macrodomain (Figure 8B) and other homologues with similarly flexible residues. The binding interactions of the fragments found in this subsite were dominated by cation-π stacking between R144 and heterocycles with some additional hydrogen bond interactions with the protein backbone, as well as interactions with Asp10 mediated by a water molecule (Fig 8C_i-vi_). This would allow merging fragments from the adenine site into this unique additional site which could lead to a higher target specificity.

**Figure 8.**
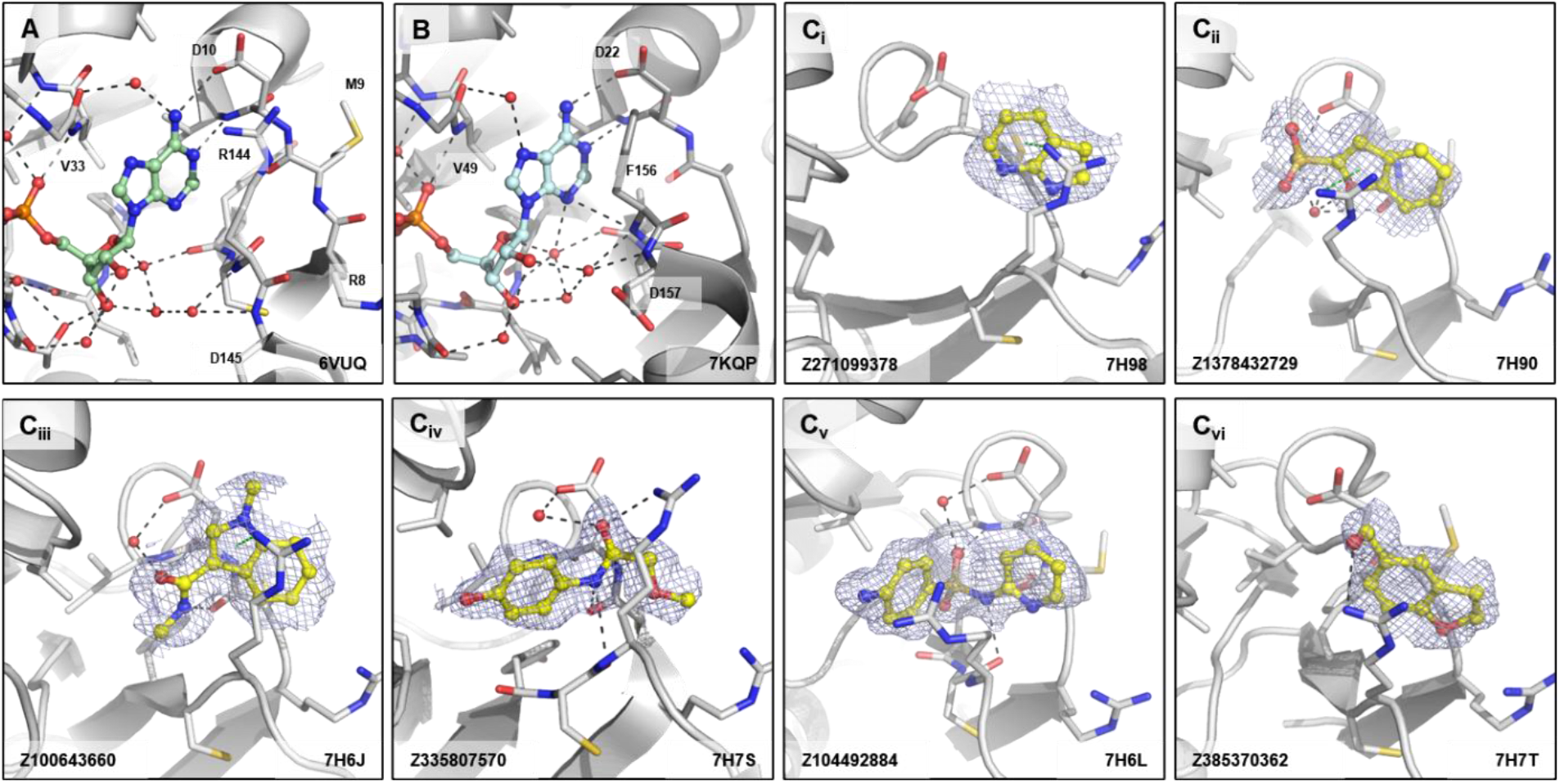
Fragments binding to the Arg144 pocket of CHIKV nsP3 macrodomain. (**A**) Stick representation of ADPr (pale green) binding to the CHIKV nsP3 macrodomain with view of Arg144 next to the adenine site. (**B**) The same view of ADPr (pale cyan) bound to the SARS-CoV-2 macrodomain which has a phenylalanine (F156) in place of the arginine residue (R144) of CHIKV. (**C**_**i-vi**_) Examples of representative fragments bound to the Arg144 pocket in CHIKV nsP3 macrodomain. The CHIKV nsP3 macrodomain structure is shown in grey. The bound fragment is shown with yellow sticks surrounded by a blue mesh generated from the PanDDA2 event map. Polar interactions and π-stacking interactions are highlighted by black and green dashes, respectively.

We, additionally, found a subset of fragments that bound in the proximity of Trp148 (‘Trp148 subsite’) of CHIKV nsP3 macrodomain (Figure 9A), which again is not present in the macrodomain of SARS-CoV-2 (Figure 9B). Fragments in this subsite were either binding to the oxyanion subsite, to Trp148 directly via π-stacking or at the backbone nitrogen of Gly112 but would expand into the solvent and not into the direction of the ADPr-binding site. Interactions were predominantly formed between a carbonyl-group and backbone nitrogen of either Arg144 in the oxyanion hole (Figure 9C_i,ii_), or of Gly112, where fragments would induce a peptide flip at Thr111 to expose the nitrogen (Figure 9C_iii-vi_). As these fragments are solvent-exposed, they could be attractive starting points for developing targeted protein degraders for the CHIKV nsP3 macrodomain.

**Figure 9.**
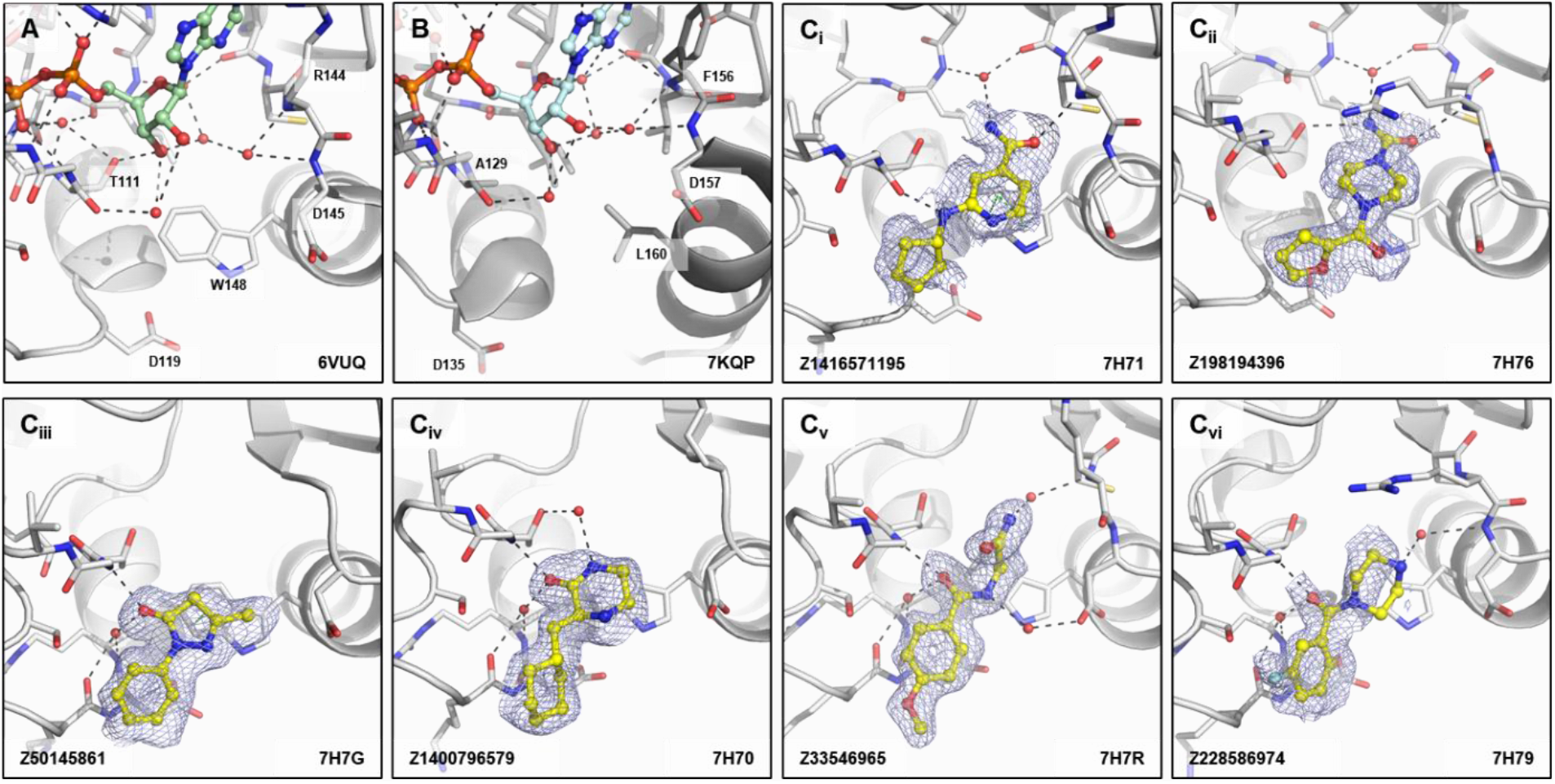
Fragments binding to Trp148 subsite of CHIKV nsP3 macrodomain. (**A**) Stick representation of ADPr (pale green) bound to the CHIKV nsP3 macrodomain with view of Trp148 below the oxyanion site. (**B**) The same view of ADPr (pale cyan) bound to the SARS-CoV-2 macrodomain which has a leucine (L160) in place of the tryptophane residue (W148) of CHIKV. (**C**_**i-vi**_) Examples of representative fragments bound to the oxyanion site in CHIKV nsP3 macrodomain. The CHIKV nsP3 macrodomain structure is shown in grey. The bound fragment is shown with yellow sticks surrounded by a blue mesh generated from the PanDDA2 event map. Polar interactions and π-stacking interactions are highlighted by black and green dashes, respectively.

### Fragment merges

The rich coverage of fragments spanning the ADPr-binding site lends itself to the design of fragment merges of fragments with overlapping pharmacophores. We manually designed three fragment merges from a combination of fragments covering the adenine and oxyanion subsites (Figure 10). We chose Z1162778919 as a starting point as it possesses high quality interactions with the oxyanion site by polar interactions and to Trp148 via π-stacking but extends upwards towards the adenine site (Figure 5B_iv_). For the adenine site, we chose three different scaffolds, representing pyrrolopyridines, amides, aminopyridine-like compounds, to merge with Z1162778919.

**Figure 10.**
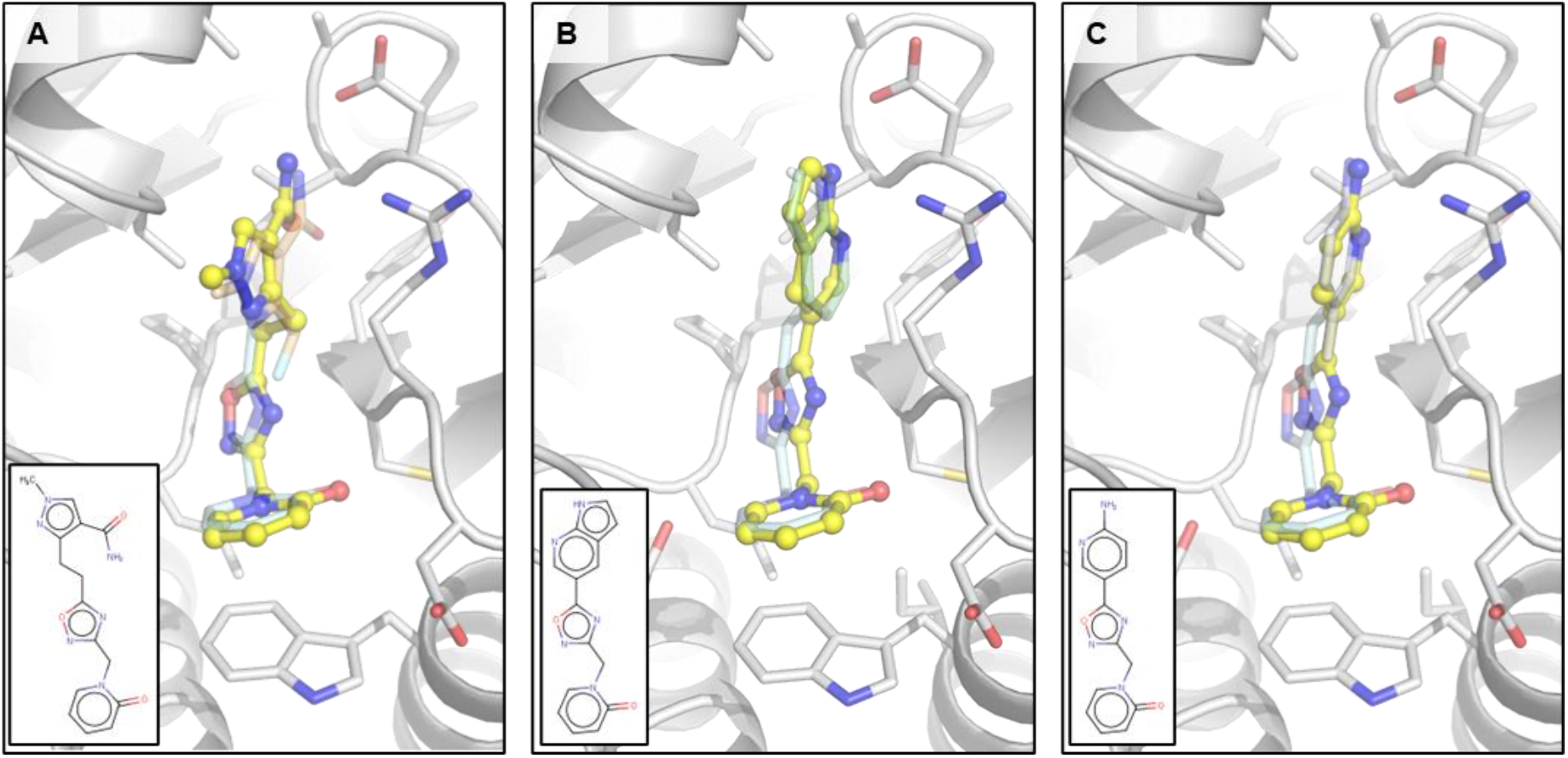
Manual fragment merges spanning the adenine and oxyanion subsite of CHIKV nsP3 macrodomain. (**A**) Fragment merge based on fragments Z1162778919 (pale cyan) in the oxyanion hole and Z1262633486 (pale orange) binding to the adenine site. (**B**) Fragment merge created by merging fragments Z1162778919 (pale cyan) in the oxyanion hole and Z271099378 (pale green) binding to the adenine site. (**C**) Fragment merge derived from fragments Z1162778919 (pale cyan) in the oxyanion hole and Z104475374 (pale blue) binding to the adenine site.

We are currently testing these designs and elaborations of them, alongside algorithmically produced merges.

## Discussion

The 109 determined hits of this crystallographic fragment screen provide novel chemical matter for the development of potent lead-like molecules that target the ADPr hydrolase activity of the CHIKV nsP3 macrodomain. Similar to the fragment screen against the SARS-CoV-2 nsp3 macrodomain^66^, the found fragments covered the ADPr-binding site with the majority of fragments occupying the adenine-binding site. The unique sub-sites we identified with our study, the Arg144 pocket and the Trp148 subsite, which do not exist in the SARS-CoV-2 macrodomain, could be additionally exploited to design an inhibitor more specific for the CHIKV nsP3 macrodomain.

Of the three available strategies for fragment progression (fragment linking, fragment merging and fragment growing^69^), we believe that fragment merging is the most rapid route to on-scale potency, which is enabled by vast structural datasets such as this one.^70,71^ Fragment merges can rapidly lead to an increase in potency by recapitulating the binding interactions initially found with the fragments. If available, these merges can be directly bought from the catalogue and tested, alternatively, usually close analogues might be available for purchase, however, in many cases, they might not fully represent the original fragment shape and interactions. To recover the full set of interactions and explore more space around the fragment merges to investigate the structure-activity relationship, multi-step synthesis can be used.

The biochemical properties and biological functions of viral macrodomains are yet to be fully understood and validated. The CHIKV nsP3 macrodomain was described to be important for viral pathogenesis and replication and its activity was shown to, additionally, counteract the host antiviral immune response.^65,72^ Targeting the CHIKV nsP3 macrodomain and, therefore, the antiviral host immune response is a differentiated approach compared to targeting viral proteins directly involved in viral replication, which comes with risks. A recent study describes the development of SARS-CoV-2 nsp3 macrodomain tool compounds which display nanomolar affinity for the macrodomain, but no antiviral activity was observed when tested in cell culture models.^73,74^ However, the sparse research that exists into alphavirus macrodomains suggests a more direct involvement of alphavirus macrodomains in viral replication compared to the SARS-CoV-2 macrodomain. A study investigating ADPr-binding and hydrolase activity of the CHIKV nsP3 macrodomain highlighted that both functions are important for virus replication. If both ADPr-binding and hydrolase activity are disrupted, the replication complex is inactive, viral RNA is not synthesised, leading to non-viable virus.^75^ Similarly, mutations in the SINV nsP3 macrodomain were found to reduce the formation of replication complexes and production of (+) and (-) strand viral RNAs.^76^ Hopefully, with the future development of inhibitors against the CHIKV nsP3 macrodomain, remaining questions regarding its biological functions can be answered.

## Conclusion

This study describes the largest, and most structurally diverse, crystallographic fragment screen of the CHIKV nsP3 macrodomain. We have rapidly made the chemical matter discovered openly available, allowing for its use in developing tool compounds and chemical probes against viral macrodomains or, more felicitously, a CHIKV or pan-alphaviral therapeutic. As part of the NIH-funded Antiviral Drug Discovery Center, READDI-AC, we are utilising the 109 fragment hits, covering the whole ADPr-binding site, as starting points for our algorithmic design of fragment merges and elaborations, which will be synthesized using our in-house chemist-assisted robotics. The subsequent designs will be analysed using our high-throughput structural biology capabilities and biophysical assays to achieve the needed structure-activity relationship information to successfully move from initial fragment hits to more lead-like molecules.

## Supporting information

Data collection and refinement statistics are summarised in the Supplementary Data File 1.

## Acknowledgements

The authors would like to acknowledge the Diamond Light Source for access to the fragment screening facility XChem, for usage of DSi-Poised and other libraries and for beamtime on beamline I04-1 under proposal LB32633. The gene for the CHIKV nsP3 macrodomain was kindly provided by A.S. de Godoy, I. Dolci and G. Oliva (São Carlos Institute of Physics, University of São Paulo).

## Author contributions

JCA carried out crystallisation optimisation, fragment screening and wrote the manuscript. JCA and DF carried out data analysis and designed manual fragment merges. CWET, BHB, RML, PGM and CW supported data collection, processing, and validation. ASG provided the construct and preliminary data. MF performed cloning, protein expression and purification, and initial crystallisation. MW and WT assisted with computational analysis and chemistry. DF and ASG reviewed the manuscript.

## Data and materials availability

All data needed to evaluate the conclusions in the paper are present in the paper and/or the Supplementary Materials. Crystallographic coordinates and structure factors for all structures have been deposited in the PDB (Protein Data Bank) (group deposition G_1002294) with the following accession codes: 7H6J, 7H6K, 7H6L, 7H6M, 7H6N, 7H6O, 7H6P, 7H6Q, 7H6R, 7H6S, 7H6T, 7H6U, 7H6V, 7H6W, 7H6X, 7H6Y, 7H6Z, 7H70, 7H71, 7H72, 7H73, 7H74, 7H75, 7H76, 7H77, 7H78, 7H79, 7H7A, 7H7B, 7H7C, 7H7D, 7H7E, 7H7F, 7H7G, 7H7H, 7H7I, 7H7J, 7H7K, 7H7L, 7H7M, 7H7N, 7H7O, 7H7P, 7H7Q, 7H7R, 7H7S, 7H7T, 7H7U, 7H7V, 7H7W, 7H7X, 7H7Y, 7H7Z, 7H80, 7H81, 7H82, 7H83, 7H84, 7H85, 7H86, 7H87, 7H88, 7H89, 7H8A, 7H8B, 7H8C, 7H8D, 7H8E, 7H8F, 7H8G, 7H8H, 7H8I, 7H8J, 7H8K, 7H8L, 7H8M, 7H8N, 7H8O, 7H8P, 7H8Q, 7H8R, 7H8S, 7H8T, 7H8U, 7H8V, 7H8W, 7H8X, 7H8Y, 7H8Z, 7H90, 7H91, 7H92, 7H93, 7H94, 7H95, 7H96, 7H97, 7H98, 7H99, 7H9A, 7H9B, 7H9C, 7H9D, 7H9E, 7H9F, 7H9G, 7H9H, 7H9I, 7H9J. Additionally, structures have been made available on Fragalysis: https://fragalysis.diamond.ac.uk/viewer/react/preview/target/CHIKV_Mac/tas/lb32633-6.

